# Sleep maintains excitatory synapse diversity in the cortex and hippocampus

**DOI:** 10.1101/2023.07.19.549645

**Authors:** Dimitra Koukaroudi, Zhen Qiu, Erik Fransén, Ragini Gokhale, Edita Bulovaite, Noboru H. Komiyama, Julie Seibt, Seth G.N. Grant

**Affiliations:** Genes to Cognition Program, Centre for Clinical Brain Sciences, University of Edinburgh, Edinburgh EH16 4SB, UK; Department of Computational Science and Technology, School of Electrical Engineering and Computer Science, KTH Royal Institute of Technology, 10044 Stockholm, Sweden; Science for Life Laboratory, KTH Royal Institute of Technology, 171 65 Solna, Sweden; Simons Initiative for the Developing Brain (SIDB), Centre for Discovery Brain Sciences, University of Edinburgh, Edinburgh EH8 9XD, UK; The Patrick Wild Centre for Research into Autism, Fragile X Syndrome & Intellectual Disabilities, Centre for Discovery Brain Sciences, University of Edinburgh, Edinburgh EH8 9XD, UK; Muir Maxwell Epilepsy Centre, University of Edinburgh, Edinburgh EH8 9XD, UK; Surrey Sleep Research Centre, School of Biosciences, University of Surrey, Guildford, Surrey, GU2 7XP, UK

## Abstract

How sleep deprivation affects cognition remains elusive. Synaptome mapping of excitatory synapses in 125 regions of the mouse brain revealed that sleep deprivation selectively reduces synapse diversity in the cortex and hippocampus. Sleep deprivation targeted specific types and subtypes of excitatory synapses while maintaining total synapse density. Altered synaptic responses to neural oscillations in a computational model suggest that sleep prevents cognitive impairments by maintaining normal brain synaptome architecture.

## Main text

Insufficient sleep is a global problem with serious consequences for cognition and mental health^1^. Synapses play a central role in many aspects of cognition, including the crucial function of memory consolidation during sleep^2^. Interference with the normal expression or function of synapse proteins is a cause of cognitive, mood and other behavioural problems in over 130 brain disorders^3^. Sleep deprivation (SD) has also been reported to alter synapse protein composition and synapse number, although with conflicting results^4-7^.

To better understand the role of sleep in regulating synapse protein composition and synapse number we have employed synaptome mapping technology to uncover the effects on SD on the mouse brain. Synaptome mapping enables highly systematic and large-scale analysis of the protein composition, protein lifetime and morphology of billions of individual synapses on a brain-wide scale^8-11^. This approach has revealed that excitatory synapses are highly diverse and can be categorised into different types and subtypes that together comprise the ‘synaptome’ of the brain^11^. These varieties of synapses are spatially distributed across all areas of the brain, forming the ‘synaptome architecture’ (SA), which changes with age and disease^8-11^. The diversity of synapse types and subtypes increases dramatically during mouse development and, after stabilising in young adults, gradually reduces with aging, with preferential preservation of synapses with the longest protein lifetimes^8,9^.

When the SA of a neuron or brain region receives patterns of neural activity it produces a spatiotemporal physiological response that is governed by the protein composition of its synapses^9-12^. Thus, changes in the SA during development, aging or disease can impact on cognition. It is not known whether the synaptome and SA of excitatory synapses change during the normal sleep-wake cycle or are affected by SD. Addressing these questions may shed light on why sleep is important for cognitive function, as well as enhance our understanding of the biological mechanisms that control synapse diversity.

Synaptome mapping of excitatory synapses in the brain utilises a line of mice that express fluorescently-labelled endogenous postsynaptic proteins PSD95 and SAP102^11^. These are scaffold proteins that assemble physically distinct multiprotein complexes^13^ and play a crucial role in synaptic transmission, synaptic plasticity and cognition^14-16^. Brain tissue sections from these mice were imaged in the mid-coronal plane at single-synapse resolution using high-throughput spinning disk microscopy with an optical resolution of approximately 270 nm. Our bespoke synaptome mapping pipeline^11^ was applied to detect and segment billions of individual synaptic puncta. We measured the density, intensity, size, shape, and colocalization parameters of each punctum. Based on these parameters, we classified each synapse into one of three main types (type 1: PSD95 only; type 2: SAP102 only; type 3: PSD95 and SAP102) and further divided them into 37 subtypes using established synapse catalogues and machine learning methods^9-11^. Finally, we spatially mapped all the measurements of individual puncta and subtypes to construct a global synaptic atlas across 125 brain regions in mice at stages throughout the normal sleep-wake cycle and after SD.

As a first step toward understanding the role of sleep in the organisation of the SA of the brain, we asked if there were any changes in the synaptome and SA during the normal circadian sleep-wake cycle. We compared three time points: the end of the active phase (zeitgeber time (ZT) 23), and at the end (ZT11) and middle (ZT6) of the resting phase. We found no differences between these time points in any of the synapse parameters or in the density of synapse types and subtypes measured in any brain area (P>0.05, Bayesian test with Benjamini-Hochberg correction) (Supplementary Fig 1, 2), indicating that the synaptome and the SA are stable throughout the normal sleep-wake cycle.

We next asked if SD induces any changes in the synaptome and SA. We compared mice after 6 hours of SD (ZT6SD) with controls that were undisturbed over the same period (ZT6). SD did not affect the density or median intensity values of PSD95-expressing and/or SAP102-expressing synapses in any of the 125 brain subregions examined (P>0.05, Bayesian test with Benjamini-Hochberg correction). However, SD did cause a decrease in the median size of synapses expressing PSD95 (P<0.05, Bayesian test with Benjamini-Hochberg correction) (Fig 1a, b) but not SAP102 (P>0.05, Bayesian test with Benjamini-Hochberg correction). The synaptome map of PSD95 puncta size showed that 99% (67/68) of cortical and 87% (20/23) of hippocampal formation (HPF) subregions were affected by SD (P<0.05, Bayesian test with Benjamini-Hochberg correction), whereas subregions in other brain areas were unaffected (P>0.05, Bayesian test with Benjamini-Hochberg correction). These results indicate that SD selectively modifies PSD95-expressing synapse types in the cortex and HPF, which are regions crucial for learning, memory and sleep consolidation^2^.

**Figure 1.**
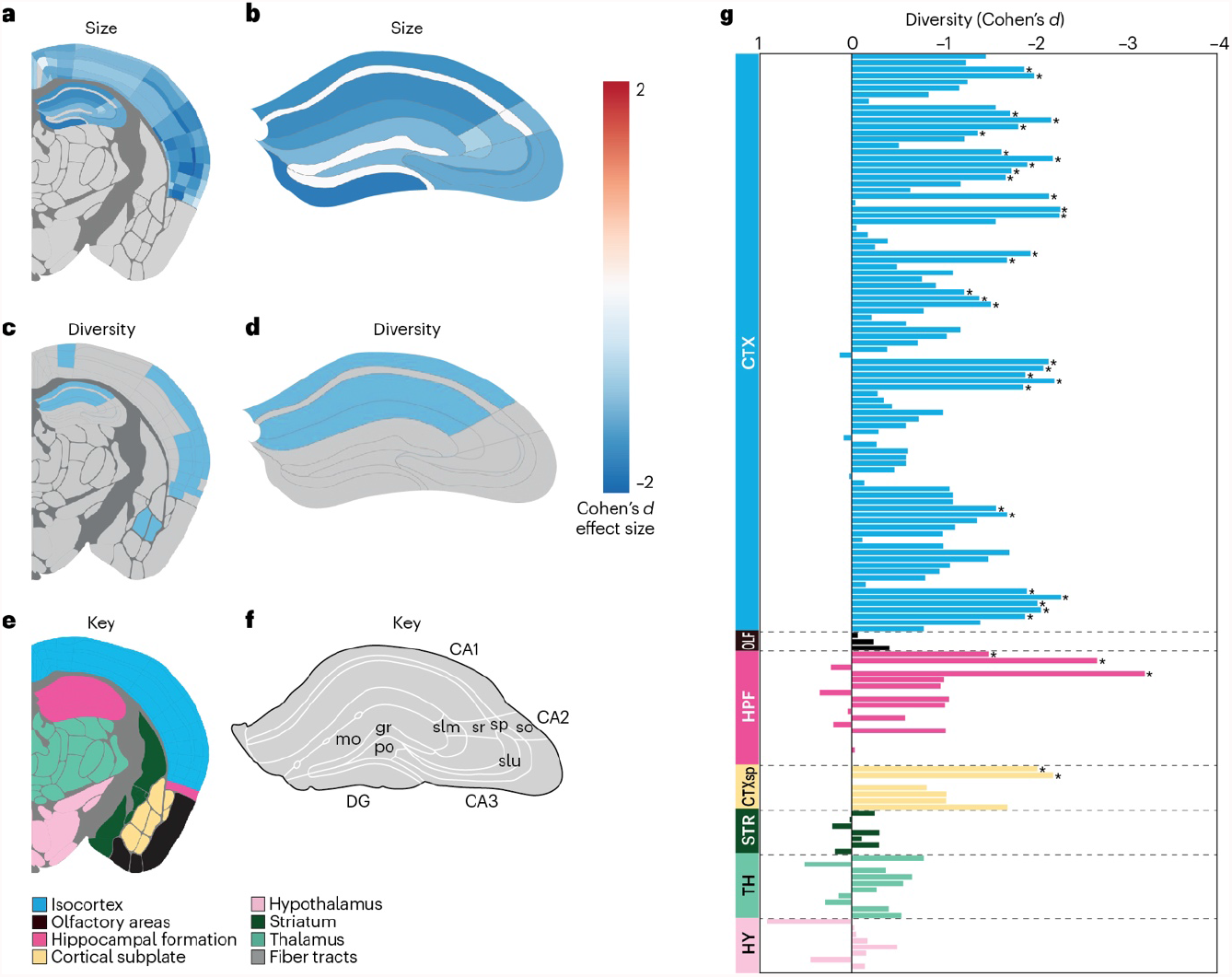
SD alters brain synaptome architecture within the cortex and HPF. SD reduces the size of PSD95-expressing synapses in cortical (a) and HPF (a,b) subregions. SD reduces synapse subtype diversity in cortical (c,g) and HPF (c,d,g) subregions. Blue regions (a-d), significant changes in Cohen’s d effect size (P<0.05, Bayesian test with Benjamini-Hochberg correction); grey regions (a-d), not significantly altered. Key for brain regions (e,f). Regions; CTX, isocortex; OLF, olfactory areas; HPF, hippocampal formation; CTXsp, cortical subplate; STR, striatum ; TH, thalamus; HY, hypothalamus.

Next, we investigated whether SD influences the high synapse diversity characteristic of the cortex and HPF regions^9,11^. By calculating the diversity of excitatory synapses based on the densities of 37 synapse subtypes, we found a significant reduction in synapse diversity across subregions of the cortex and the HPF in response to SD (Fig 1c, d, g). To better understand how this reduction in synapse diversity was reflected among the 37 synapse subtypes, we created a heatmap of the density change (Cohen’s d) for each subtype in cortex and HPF (Supplementary Fig 3). This shows that the density of some subtypes was consistently reduced across these regions whereas others were simultaneously increased, indicating that SD drives changes in the same subtypes in different neuron types. Furthermore, because there is no net change in synapse density, the subtypes that are reduced in density may be transforming their identity to those subtypes that are increased in density.

In previous studies, we discovered that certain ‘short protein lifetime’ (SPL) subtypes of excitatory synapses exhibit faster turnover rates for PSD95 than others^8^. These SPL subtypes were involved in synaptic adaptations to mutations and aging^8,10^. By contrast, other ‘long protein lifetime’ (LPL) subtypes, with a slower rate of PSD95 turnover, were selectively preserved in older individuals and suggested to play a role in long-term memory storage^8,9^. Based on these findings, we hypothesized that the synapse subtypes most likely to undergo changes in response to SD would be those previously demonstrated under aging and mutation challenges to exhibit higher adaptability. Supporting our hypotheses, a heatmap representing the change in density of subtypes, ranked by their protein lifetime, across all regions of the cortex and HPF indicates that subtypes with longer protein lifetimes generally increased, whereas those with shorter lifetimes mostly decreased (Fig 2a). Additionally, the relationship between the lifetime of PSD95 protein and the change in density (Cohen’s d) of each subtype in the cortex reveals that synapses with shorter protein lifetimes experienced a decrease, whereas those with longer protein lifetimes showed an increase (Fig 2b). To further examine this, we compared the density changes between subtypes with the longest and shortest protein lifetimes. This revealed that the subtypes with longer lifetimes exhibited significantly greater increases than those with shorter lifetimes in various cortical subregions, olfactory regions, cortical subplate, and select HPF subregions (P<0.05, Bayesian test with Benjamini-Hochberg correction) (Fig 2c). In summary, our findings suggest that SD leads to a reduction in the diversity of excitatory synapses overall, primarily driven by a decrease in the number of synapse subtypes with short protein lifetimes and an increase in those with long protein lifetimes.

**Figure 2.**
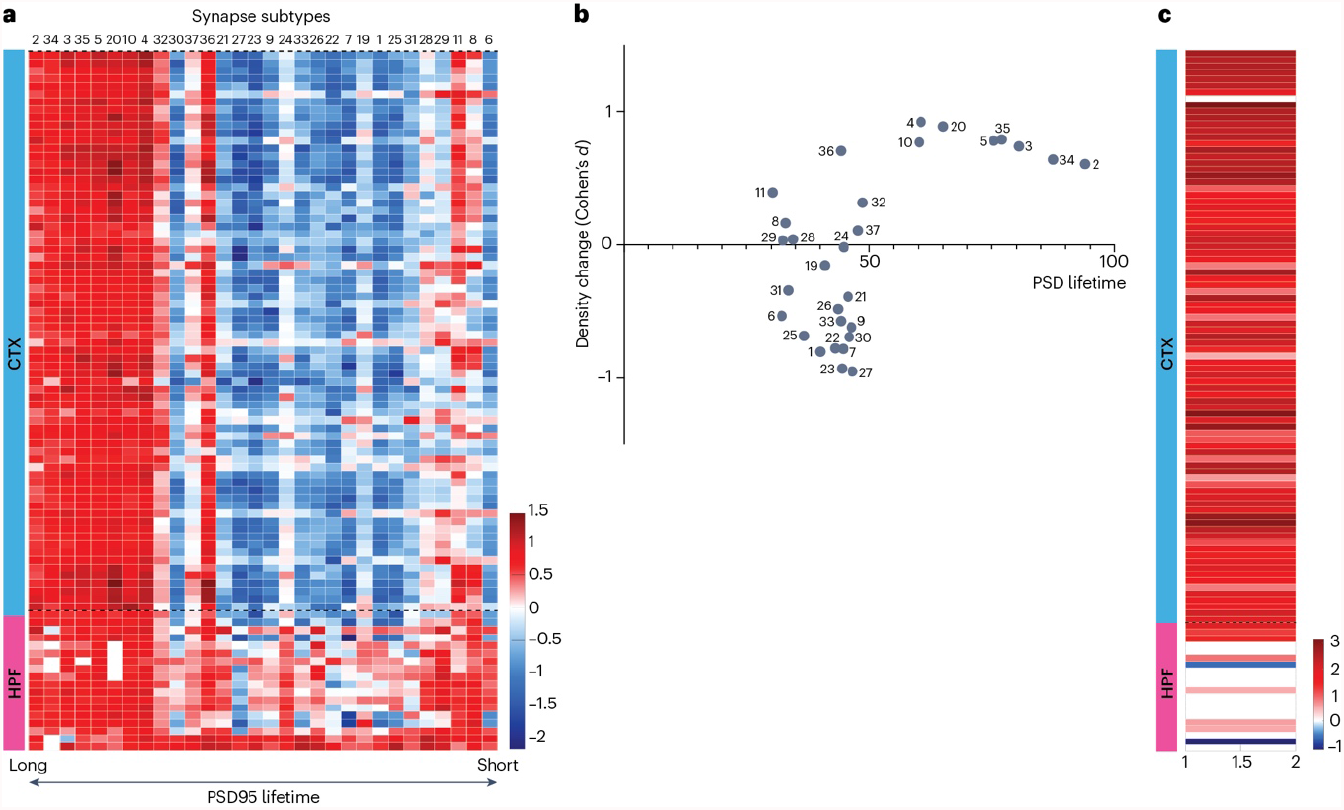
SD differentially impacts synapse subtypes. (a) Heatmap of SD-induced changes in the density (Cohen’s d) of synapse subtypes in cortex (CTX) and HPF ranked from longest to shortest PSD95 lifetime^8^. For a heatmap showing significant changes corrected for multiple testing see Supplementary Fig 4. (b) SD-induced synapse subtype density changes in the cortex (average of all subregions) plotted against PSD95 lifetime (normalized percentage)^8^. (c) Comparison of the density changes of six LPL (2, 3, 5, 20, 34, 35) with six SPL (6, 8, 11, 28, 29, 31) subtypes after SD. Red signifies subregions with greater change (Cohen’s d) in LPL than SPL synapses, whereas blue signifies subregions with greater change in SPL than LPL synapses; white subregions show no significant differences.

Distinct patterns of neural activity are recognized as a defining characteristic of sleep and wakefulness, and it is widely believed that these patterns contribute to the process of memory encoding^8^. The SA plays a crucial role in converting patterns of neuronal activity into a spatiotemporal output^11,12^ and this process can be modified when there are changes in the SA^9-11^. To investigate whether SD-induced changes in the SA of the CA1 stratum radiatum (CA1sr) in the HPF could affect responses to activity patterns related to sleep, wakefulness, and memory encoding, we employed a well-established computational model^9-11^ (Fig 3a). When stimulating the CA1sr with five distinct activity patterns (gamma train, gamma burst, theta train, theta burst, sharp-wave ripple), each pattern had a different effect on SD versus control (P=0.01, paired Kolmogorov-Smirnov test, N=121; Benjamini-Hochberg corrected, N=5; Cohen’s d > 1.9) (Fig 3b). Notably, the most pronounced effects were observed in spatial responses for theta burst and gamma train patterns, while the intensity of responses was primarily influenced by the sharp-wave ripple pattern (Fig 3b, c).

**Figure 3.**
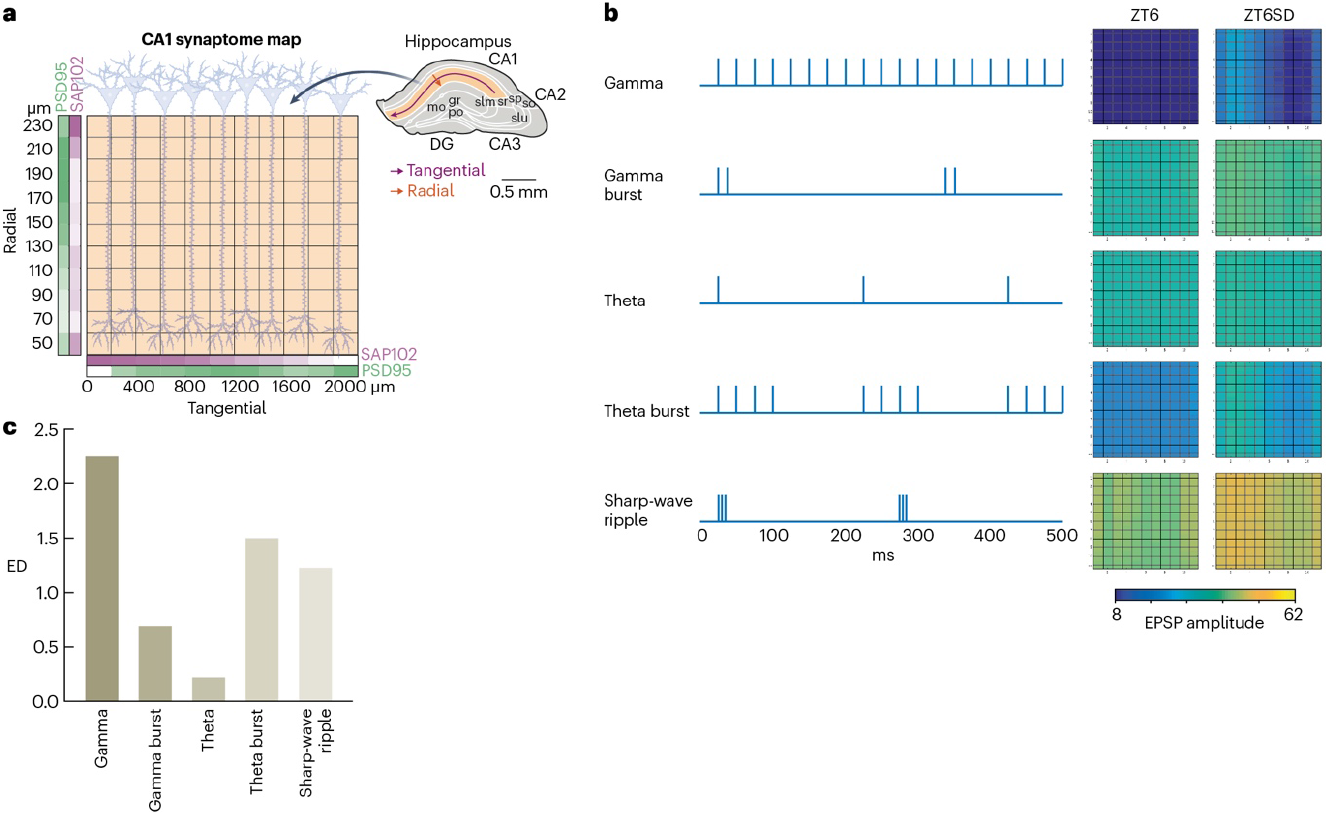
Computational modelling of physiological responses in the CA1sr to patterns of activity after SD. (a) The model simulates a 2D (11 × 11) array of synapses (boxes) expressing PSD95 and SAP102 measured along the radial and tangential axes of the CA1sr^10,11^. (b) Five patterns of neuronal activity were used for CA1sr stimulation in the computational model. The summed excitatory postsynaptic potential (EPSP) response amplitudes in the ZT6 and ZT6SD groups were quantified (colour bar, arbitrary units). (C) Comparison of the extent of disruption caused by SD for each of the five patterns of activity. ED, Euclidian distance.

Our findings highlight the significance of sleep in preserving synaptome diversity in the cortex and HPF, which are brain regions specialized for memory and higher cognitive functions. Lifespan studies of the SA identified a correlation between synaptome diversity and the acquisition of complex behaviours during development and their impairments with aging^9^. By preserving synaptome diversity, sleep may contribute to optimizing cognitive function on a daily basis.

Within just 6 hours of SD there was a shift in the populations of synapse subtypes toward LPL synapses. A similar shift occurs in the aging brain where LPL synapses are enriched^8,9^. The slower rate of protein turnover in LPL synapses may render the sleep deprived and aged brain less adaptable, potentially impairing learning and repair. Consistent with this, the repair of the synaptome architecture in *Pax6* mice during development is primarily mediated by SPL synapses^10^. Together these observations suggest that synapse subtypes with rapid proteostasis play a vital role in the brain’s adaptive responses to environmental and genetic perturbations.

Our current understanding of the mechanisms behind the impairments in memory consolidation induced by SD is limited. Our study suggests the synaptome and SA may play a role from several perspectives. Firstly, we found that SD specifically affects the SA of the HPF and cortex, regions crucial for memory consolidation during sleep^17,18^. Secondly, as a consequence of the altered SA the electrophysiological responses in the CA1sr to theta burst and gamma train patterns were modified. These changes could impact the encoding and learning phases in the active state of the animal, ultimately affecting memory consolidation during sleep^19,20^. Furthermore, modifications in sharp-wave ripple activity could contribute further to memory impairments associated with SD by influencing replay activity supported by these oscillations during periods of quiet waking (when sleep-like patterns are observed in the HPF). Superimposed on these electrophysiological functions is the role of protein turnover, which Francis Crick highlighted almost four decades ago^21^ as a crucial determinant of memory consolidation. The imbalance between synapses with long and short protein lifetimes caused by SD might itself interfere with consolidation and even synergise with the altered electrophysiological responses. The modification of the SA of the brain offers a new framework for understanding the synaptic basis of SD and suggests that interventions targeting specific subtypes of synapses is an avenue that could potentially prevent or reverse cognitive impairments associated with SD and aging.

## Material and Methods

### Animals

Animal procedures were performed in accordance with UK Home Office regulations and approved by Edinburgh University Director of Biological Services. Generation of the *Psd95*^eGFP/eGFP^;*Sap102*^mKO2/mKO2^ knock-in mouse line using C57BL/6 J mice and its characterisation have been described^11^. The same study^11^ was used to estimate the sample size needed, leveraging a t-test analysis. Adult 3-month-old mice were used for this study. Mice were transferred from their home cage to a new, conventional, enrichment-free cage, where they were single-housed for a 3-day habituation period (days 0-2) and for the day of the experiment (day 3). The animals had ad libitum access to water and food and were kept in a quiet room of ∼22°C and 12-hour light:dark cycle (lights on 7 am, lights off 7 pm). For the purposes of masking the experimental groups to the experimenter during processing of samples and analysis, each animal received a randomized ID number which was maintained throughout tissue processing and imaging.

### Circadian sleep-wake cycle study

A group of 30 animals was split into three equivalent groups of 10 animals, each comprising 5 males and 5 females. The mice were undisturbed during days 0-2 and their brain tissue was collected at ZT6, ZT11 or ZT23 on day 3 based on their assigned group.

### Sleep deprivation study

A group of 18 animals was split into two groups: the ZT6 group (5 males and 3 females) and the ZT6SD group (6 males and 4 females). All mice were handled daily for 10 minutes by the experimenter between 07:30 am and 08:00 am during habituation days 0-2. On day 3, the mice were either left undisturbed (ZT6 group) or were kept awake by gentle handling (ZT6SD group), which comprised gentle auditory and tactile stimulation (tapping on the cage and touching the animals with a brush) when the animals were visually observed to have fallen asleep. The brain tissue of animals of both groups was collected 6 hours after lights-on (time point ZT6). The experimenter was present in the room during the 6 hours of the experiment on day 3 for both groups.

### Tissue collection and sectioning

At the collection time points mice were anaesthetised by intraperitoneal injection of a lethal dose of 0.1 ml 20% pentobarbital (Euthatal, Merial Animal Health). After complete anesthesia, 10 ml phosphate-buffered saline (PBS; Oxoid, Basingstoke, UK) were used per animal for cardiac perfusion, followed by 10 ml 4% (v/v) paraformaldehyde (PFA; Alfa Aesar, Heysham, UK) for fixation, both solutions at 4°C. Whole brains were dissected out and immediately postfixed at 4°C in 5 ml 4% PFA for 4 hours, before being transferred to 5 ml 30% (w/v) sucrose (VWR Chemicals, Lutterworth, UK) in 1×PBS at 4°C for ∼72 hours. In preparation for embedding, brains were kept for 1 hour in a 1:1 solution of 30% sucrose and Optimal Cutting Temperature Medium (OCT, VWR International) at 4°C. Finally, brains were placed in plastic cubic molds (Sigma-Aldrich) containing OCT for embedding, and were immediately frozen in a container with ∼10 ml isopentane (Sigma-Aldrich) placed in liquid nitrogen. After completion of embedding, the tissue was stored at -80°C. Each brain was sectioned in the coronal -1.9 mm bregma level at 18 μm thickness using a cryostat (NX70, Thermo Fisher Scientific, Gloucester, UK), and placed on glass SuperFrost Plus microscopy slides (Thermo Scientific).

### Tissue preparation

In preparation for imaging, the brain tissue slices were rinsed with ice-cold 100 μl PBS for 5 min. Excess PBS was then removed with a Kimwipe tissue, and 12 μl Mowiol mounting medium (96 g glycerol (Sigma-Aldrich, BioXtra >99%), 38.4 g Mowiol (Calbiochem), 192 ml 0.2 M Tris buffer (pH 8.5), 96 ml milliQ water (18.2 MΩ)) with 2.5% DABCO (1,4-diazabicyclo[2.2.2]octane; Sigma-Aldrich) was added to each section for optimal imaging without absorption, autofluorescence or light scattering, and for the prevention of photobleaching. Finally, each slice was covered with a 13 mm diameter, 1.5 mm thick round glass coverslip (VWR International) and imaged the following day.

### Spinning-disk confocal microscopy

Image capture employed an Andor Revolution XDi spinning-disk microscopy system equipped with a Yokogawa CSU-X1 50 μm pinhole spinning disk, an Olympus uPlanSAPO 100X oil-immersion lens (NA 1.4), and an Andor iXon Ultra monochrome back-illuminated EMCCD camera capturing images with 16-bit depth and 512 × 512 pixels (pixel size of 0.084 μm). Frame averaging of 2 and 250 EMCCD gain was applied to all synaptome mapping scans. The imaging settings for each fluorophore-tagged protein in the synaptome mapping experiments were: PSD95eGFP 488 nm, QUAD emission filter, exposure time 0.07 s, power 20%; and for SAP102mKO2 561 nm, QUAD emission filter, exposure time 0.1 s, power 40%. A single mosaic grid was used to cover each entire brain section with an adaptive z-focus set up by the user to follow the unevenness of the tissue using Andor iQ2 software.

### Synaptome mapping pipeline

The synaptome mapping (SynMap) technique^11^ was established and standardized to systematically map individual PSD95eGFP puncta across the entire brain. Using deep learning methods developed in house, SynMap comprises a sequence of automated image analysis procedures that includes puncta detection, colocalization, classification, and map reconstruction, among others. To define the anatomical regions within SynMap, manual delineation was employed, using the Allen Reference Atlas (http://mouse.brain-map.org/) as a guiding resource for identifying the boundaries of distinct anatomical areas.

### Cohen’s d formula

Cohen’s d effect size was calculated as per Cohen^22^ as follows:

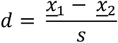

Where <u>*x*</u> is the mean of one of the groups, where *s* is the pooled standard deviation as

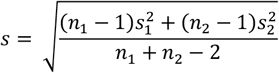

where the variance (*s*^2^) of one of the groups as

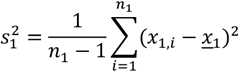

### Bayesian analysis

Bayesian estimation^23^ as used previously was applied to provide a more objective statistical test of the estimate the significance of the effects of SD on synaptome maps, including basic PSD95 parameters, subtype density and diversity.

A two-stage Bayesian estimation approach was used to examine the differential effects of SD on SPL and LPL subtypes. In the first stage, a probability distribution of effect size for both SPL and LPL regarding SD was generated. In the second stage, an additional Bayesian estimation was applied to the previously estimated distribution to determine the effects of SD on SPL or LPL subtypes. This two-stage testing method was implemented for each subregion to generate significant *P*-values and median Cohen’s d values. Lastly, Benjamini-Hochberg corrections were applied to compute the corrected *P*-values across all brain regions.

### Computational modeling of synaptic responses

Computational modeling of synaptic temporal responses was based on our previously described model^9-11^ representing physiology in the hippocampus, briefly outlined below. Here we modified this model to include the effects of timepoint and SD state. These synaptic models simulate changes in synapse temporal responses, EPSP amplitudes, short-term plasticity and temporal summation based on observations from neurons where PSD95 and SAP102 expression is altered^14,15,24-26^. For a description of the modeling of how the spatial differences in PSD95 and SAP102 affects individual synaptic time dynamics, see^10^.

#### Synaptic scaling representing timepoint and SD state

As described in previous work^9,11^, the size of PSD95 and SAP102 synapses along the radial and tangential directions of the hippocampus (Fig. 3) is derived from the fluorescence intensity measurement of individual synaptic puncta and represented by color intensity; PSD95 (green) and SAP102 (magenta). These size values were used to scale the synaptic properties of the computational model to represent differences in animal group (timepoint and SD state). To model synaptic physiology corresponding to 3 month old animals in control and sleep deprived animals, differences in the size of PSD95 and SAP102 along radial as well as tangential directions of hippocampus were computed as outlined below.

The hippocampus was delineated into four tangential subregions (CA1, CA2, CA3 and dentate gyrus) and four radial layers (for CA1-3 stratum lacunosum-moleculare, stratum radiatum, stratum pyramidale and stratum oriens and for dentate gyrus the superior molecular layer, polymorphic layer, inferior granular layer and inferior molecular layer), for each of the animal groups. Next, we computed the geometric mean over individuals (N=8 for ZT6 and N=10 for the other groups) of PSD95 and SAP102 puncta size. Then, for each protein, we normalized data according to the following. We computed the directional gradient (largest minus smallest value) along each direction (radial or tangential) for each animal group and from this identified the minimum and the maximum span over all animal groups. Normalization was done by subtracting the minimum span and dividing by that maximun span. Thus, for radial and tangential size values of PSD95 and SAP102 puncta, the expression was compared over all animal groups. This relative size was used to scale the spatial distribution of synaptic values used in previous work^11^. This scaling thus allows for a comparison of animals at different daily time points and SD.

## Acknowledgements

B. Notman, T. Wong, E. Sigfridsson, B. Koniaris for technical assistance, C. Davey for editing, D. Maizels for artwork.

## Funding

The European Research Council (ERC) under the European Union’s Horizon 2020 Research and Innovation Programme (695568 SYNNOVATE) DK, EB, ZQ, SGNG. Wellcome Technology Development Grant (202932/Z/16/Z) RG, ZQ, SGNG. Simons Foundation Autism Research Initiative (529085) NHK, SGNG. Wellcome Trust (209099/Z/17/Z) JS. Leverhulme Trust (RPG-2020-340) JS. DK was part funded by a Principal’s Career Development PhD Scholarship from The University of Edinburgh. For the purpose of open access, the author has applied a CC-BY public copyright license to any author accepted manuscript version arising from this submission.

## Author contributions

DK planned the experiments, collected, imaged and analyzed brain samples, performed image and data analyses. ZQ developed software and performed image and data analyses. EF analyzed data and performed computational modeling of neural activity. RG performed data handling and generating brain maps. EB provided data on synapse protein turnover. NHK and JS provided supervision. SGNG supervised the project and wrote the manuscript.

---

**Supplementary Figure 1.**
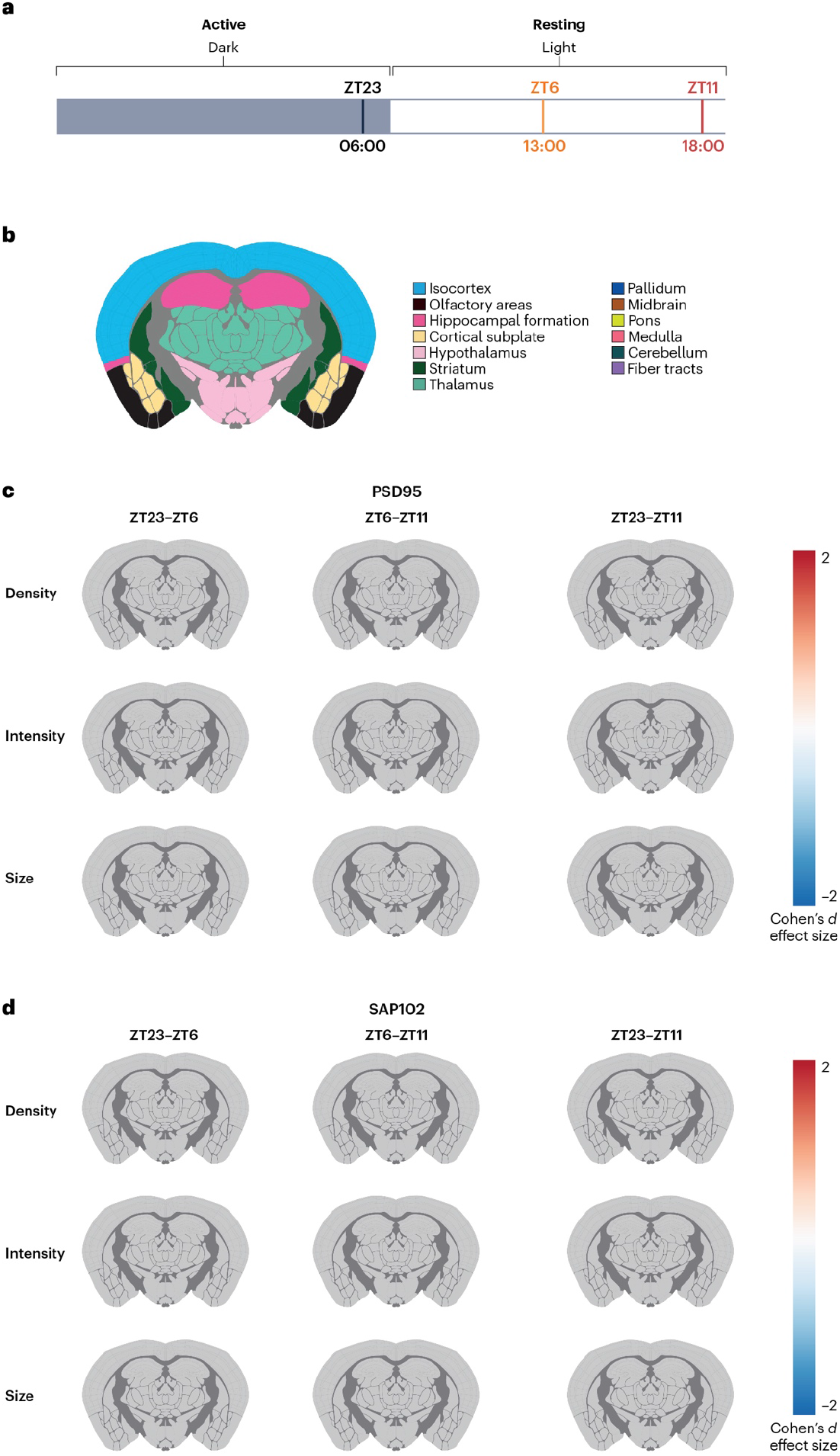
The synaptome architecture of PSD95 and SAP102 puncta does not change in the circadian cycle. (a) Schematic representation of light/dark cycle, with zeitgeiber time (ZT) showing when mice were sampled (ZT23, ZT6, ZT11) and corresponding 24-hour clock (06:00, 13:00, 18:00). (b) Key for brain regions shown in maps (c,d). (c) Synaptome maps of PSD95 puncta density (top row), intensity (middle row), size (bottom row) changes for periods ZT23-ZT6 (left column), ZT6-ZT11 (middle column), ZT23-ZT11 (right column) in brain regions shows no significant changes (grey; P>0.05, Bayesian test with Benjamini-Hochberg correction). (d) Synaptome maps of SAP102 puncta density (top row), intensity (middle row), size (bottom row) changes for periods ZT23-ZT6 (left column), ZT6-ZT11 (middle column), ZT23-ZT11 (right column) in brain regions shows no significant changes (grey; P>0.05, Bayesian test with Benjamini-Hochberg correction).

**Supplementary Figure 2.**
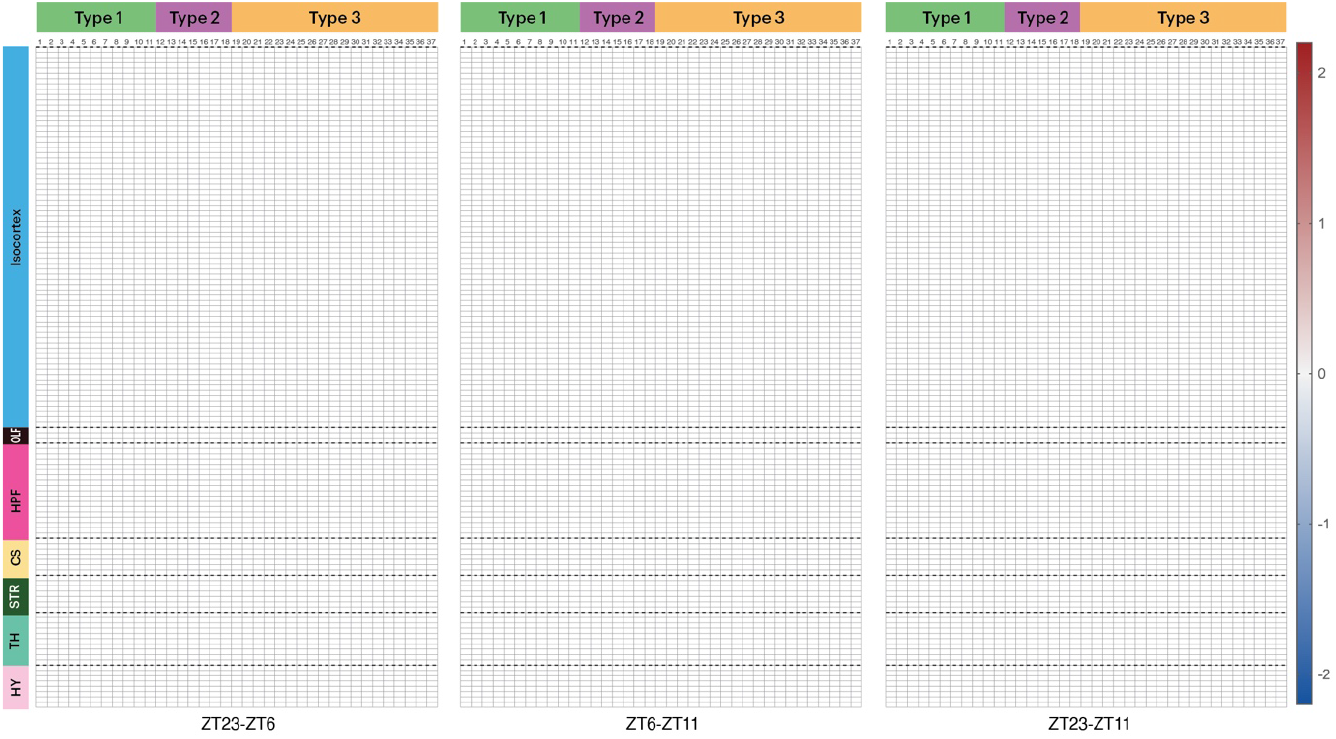
The synaptome architecture of Type 1, Type 2 and Type 3 synapse subtypes does not change in the circadian cycle. Heatmaps of SD-induced changes in the density (Cohen’s d) of synapse subtypes in brain regions for periods ZT23-ZT6 (left panel), ZT6-ZT11 (middle panel), ZT23-ZT11 (right panel). All datapoints show P>0.05, Bayesian test with Benjamini-Hochberg correction. Regions; isocortex; OLF, olfactory areas; HPF, hippocampal formation; CS, cortical subplate; STR, striatum; TH, thalamus; HY, hypothalamus.

**Supplementary Figure 3.**
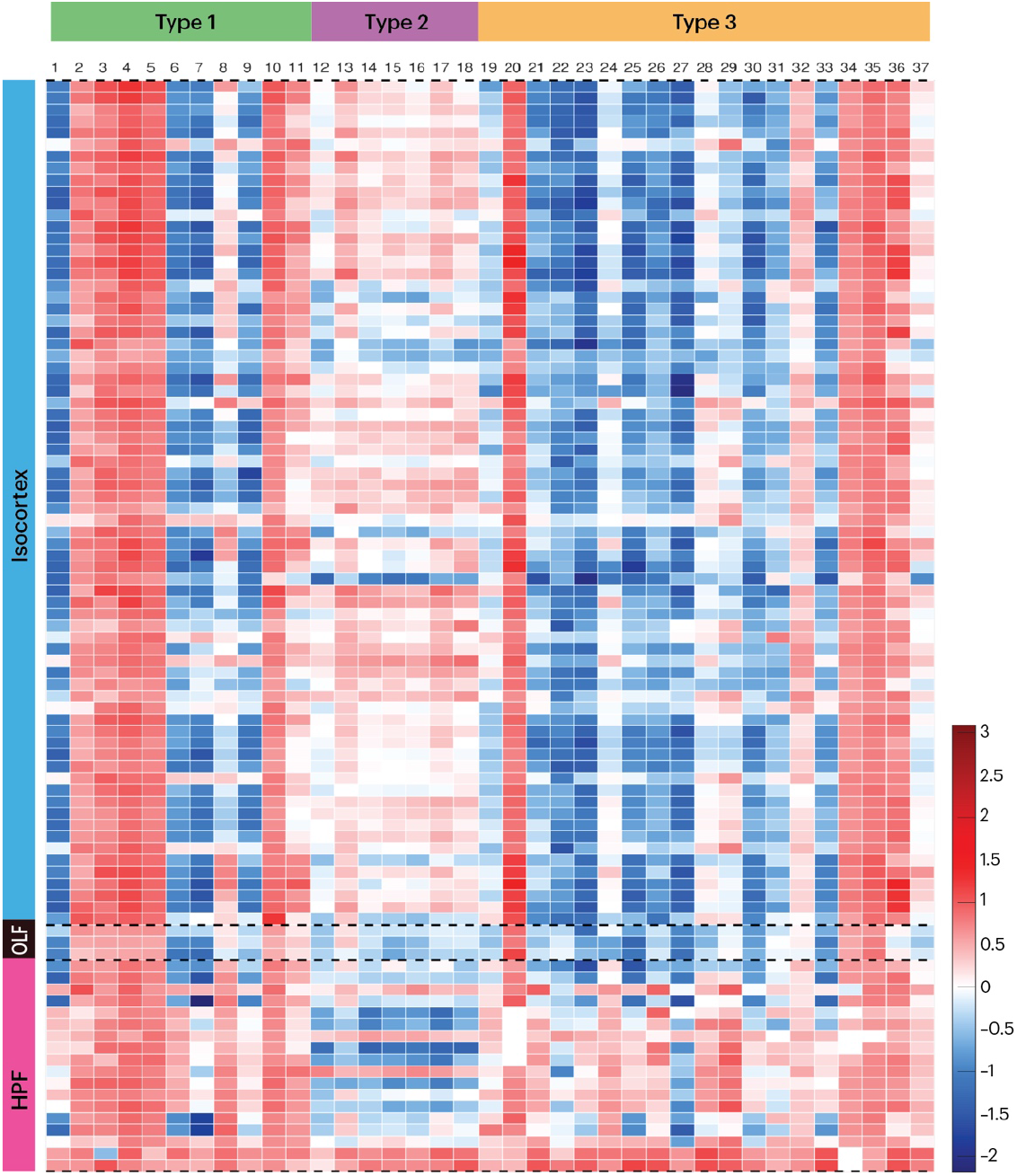
SD differentially impacts synapse subtypes. Heatmap of SD-induced changes in the density (Cohen’s d) of synapse types and subtypes. Uncorrected. Significant changes are shown in Supplementary Fig 4. Regions: isocortex; OLF, olfactory regions; HPF, hippocampal formation.

**Supplementary Figure 4.**
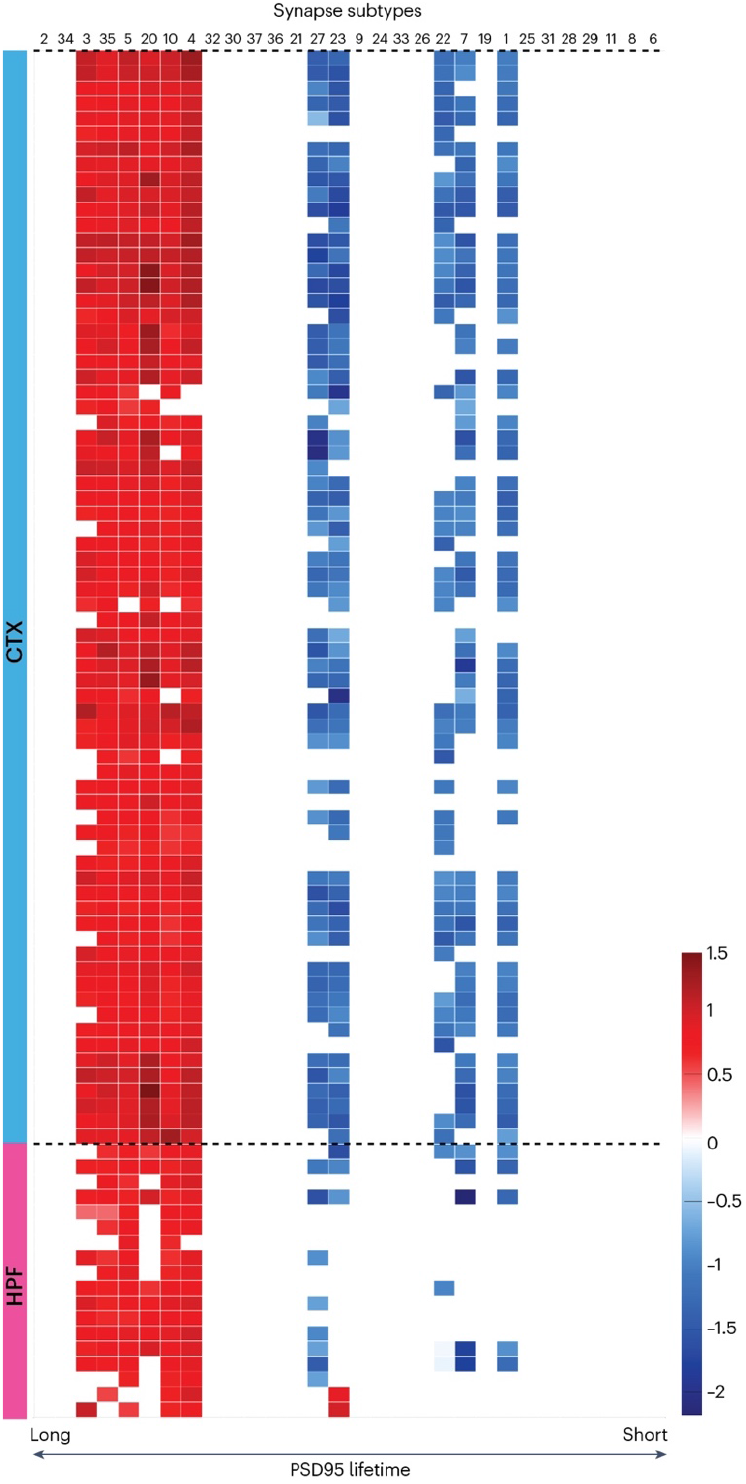
SD differentially impacts synapse subtypes with long and short protein lifetimes. Heatmap of SD-induced changes in the density (Cohen’s d) of synapse subtypes ranked from longest to shortest PSD95 lifetime^8^. Significant changes are shown (P<0.05, Bayesian test with Benjamini-Hochberg correction).

## Notes

### Competing Interest Statement

The authors have declared no competing interest.

